# Anomalous Scaling of Gene Expression in Confined Cell-Free Reactions

**DOI:** 10.1101/251306

**Authors:** Ryota Sakamoto, Vincent Noireaux, Yusuke T. Maeda

## Abstract

Cellular surface breaks the symmetry of molecular diffusion across membrane. Here, we study how steric interactions between the surface and the bulk of cell-sized emulsion droplets alters gene expression emulated by a cell-free transcription/translation (TXTL) system. The concentration of synthesized reporter proteins in droplets of radius *R* shows an anomalous geometric scaling of *R*^4^ different from the expected normal size-dependence of *R*^3^. Given that TXTL becomes less efficient at thin surface layer, a mathematical model explains anomalous size-dependence found in experiment. The surface of cell-sized compartment thus plays a regulatory role for cell-free gene expression.

## Introduction

As expressed by W. Pauli “God made the bulk, surfaces were invented by the devil.”, the surface has remarkable properties due to its anisotropic nature, especially in microscopic systems. The atoms or molecules close to the surface have three different types of interaction: with the inner phase, with the outer phase and with the neighbors on the surface. The fact that broken symmetry in surface alters the material properties has been widely accepted in condensed matter physics, from semiconductors^1^ to colloidal transport^2^. In biological systems, the smallest functional and structural unit, which has a functional bulk space enclosed in a thin interface, is a living cell. Typical size of a cell ranges from 1 to 100 μm^3^. Consequently, the surface area to volume ratio (S/V ratio) plays an important role for cellular physiology. On the one hand, lasting exposure to the environment enhances the uptake nutrient or water with higher efficiency^4^.

On the other hand, the surface of cells faces the inner cytosolic space in which genetic information stored in DNA can be expressed through transcription (TX) and translation (TL)^5,6^. Cell-free expression systems (CFES), such as the PURE system^7,8^ or TXTL^9,10,11,12,13^, recapitulate gene expression in vitro. CFES have become ideal platforms to construct cellular functions in isolation through the expression of synthetic gene circuits. TXTL, used in this work, is based on a cytoplasmic extract prepared from *E. coli* that contains the molecular machineries for efficient transcription and translation. TXTL is increasingly used to engineer models of synthetic cells^14,15,16,17^. CFES reactions of both PURE system and TXTL are also encapsulated into cell-sized emulsion droplets or liposomes programmed with gene circuits for a specific cellular function^18,19,20,21^. At the fundamental level, however, a lot remains to be understood on how gene expression is altered in such microscopic compartments. While the variability in the kinetics of cell-free gene expression in confined environments has been extensively discussed^22,23,24,25,26,27,28,29,30^, the functional interplay between the surface, TXTL reactions, and intracellular metabolism remains elusive and poorly characterized.

In this study, we demonstrate that cell-sized compartments alter the geometric scaling law of gene expression. We derive a model to show that the geometric scaling between produced protein and the radius of a compartment *R* reflects surface-induced gene regulation: when surface represses or activates gene expression, the protein amount scales with *R*^4^ or *R*^2^ respectively. Experimentally, we show that the concentration of a synthesized reporter protein increases in a quartic manner as a function of droplet radius when TXTL is executed in cell-sized emulsion droplets. This anomalous geometric scaling supports interfacial repression of TXTL. Our findings provide new insight for minimal bioreactor out of equilibrium.

## Results

### Cell-free gene expression in droplets

The TXTL system used to perform CFGE in emulsion droplets (Fig. 1) was described previously^11,12^. The TXTL system contains all the necessary proteins and metabolites for CFGE. The plasmid P_70*a*_-deGFP (3.2 kbp, used at 16 nM)^11^ was used to express the reporter protein deGFP (green fluorescent protein). Rhodamine-dextran (Sigma-Aldrich) was also used as a red fluorescent marker to determine the droplet volumes (∝ *R*^3^). The emulsion droplets were prepared using the phospholipid 18:1 (∆9-Cis) phosphatidyl-choline (DOPC, Avanti Polar Lipids) as surfactant in mineral oil (Sigma-Aldrich). The DOPC lipid was dissolved in the oil at 0.1% (w/v) so as to make reaction-in-oil droplet (DOPC droplet) coated by a lipid monolayer. DOPC droplets encapsulating TXTL reaction with DNA were prepared by adding 30*μ*L DOPC oil to 12*μ*L TXTL reaction, then emulsified by tapping on the tube. The size of DOPC droplet was set at its radius *R* ≥ 5*μ*m. The average number of plasmid DNA is 5000 molecules within smallest droplet of 0.5 pL. The 10*μ*L of the emulsion was placed onto a slide glass with polydimethyl-siloxane coat and enclosed by frame seal chamber with cover slip. Time-lapse of GFP fluorescence kinetics were recorded at time interval of 5 min for 15 hours by epi-fluorescence (IX73, Olympus microscope) with a cooled CMOS camera (Neo5.5, Andor). The size of droplets was analyzed by a custom-made MATLAB image processing. The temperature was kept at 31°C using a custom-made hot water bath attached to the microscope stage.

**Figure 1.**
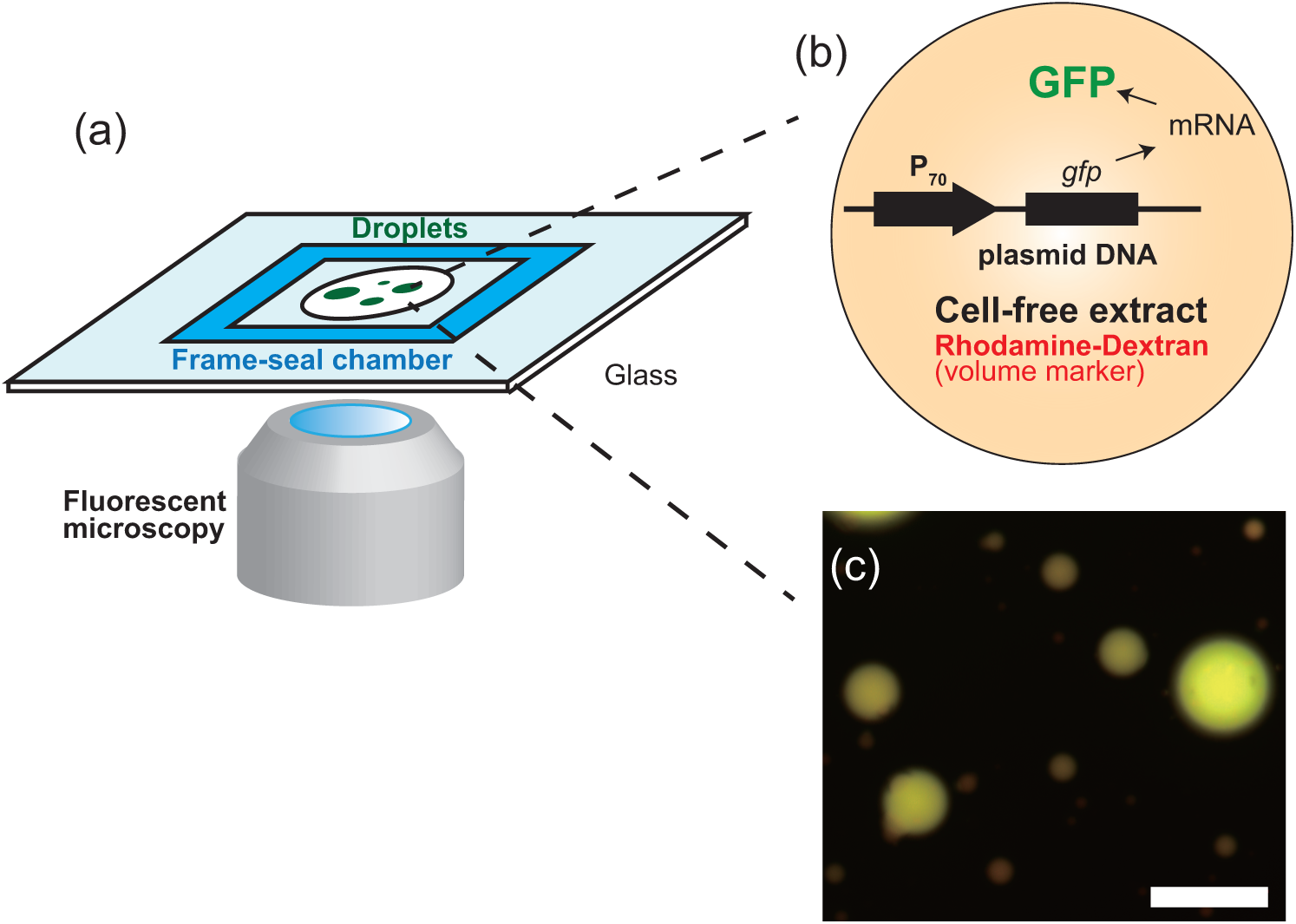
(a) Schematic illustration of experimental setup. Emulsion droplets containing the TXTL extract and plasmid DNA are recorded in multi-color fluorescent microscopy. (b and c) The product of CFGE, deGFP, was observed in green fluorescent channel while Rhodamine-dextran as volume marker was visualized in red-fluorescent channel. Scale bar: 100 *μ*m.

Poly-dispersed droplets, with diameter ranging from few micron to ~100 *μ*m, were prepared to characterize CFGE of a reporter gene in confined environment. Expression of the reporter gene *degfp* was driven by the promoter P_70_, within the emulsion droplets (Fig. 1(b))^11^. The overall system recapitulates gene expression in vitro in physiological conditions so as to mimick conditions found in living cells^14^. Fig. 1(c) shows DOPC droplets containing the synthesized deGFP after 15 h of incubation with Rhodamine-dextran. To make sure the reliable steady state (end-point) of CFGE, we measured the kinetic time-course, how long it continues to synthesize proteins and then stops. As shown in Fig. 2, the time-course of deGFP fluorescence in emulsion exhibited monotonic increase, each curve corresponds to single droplets varied in size. The fluorescence level has reached plateau after 15 hours, indicating that CFES reached chemical equilibrium where all the reactions stop.

**Figure 2.**
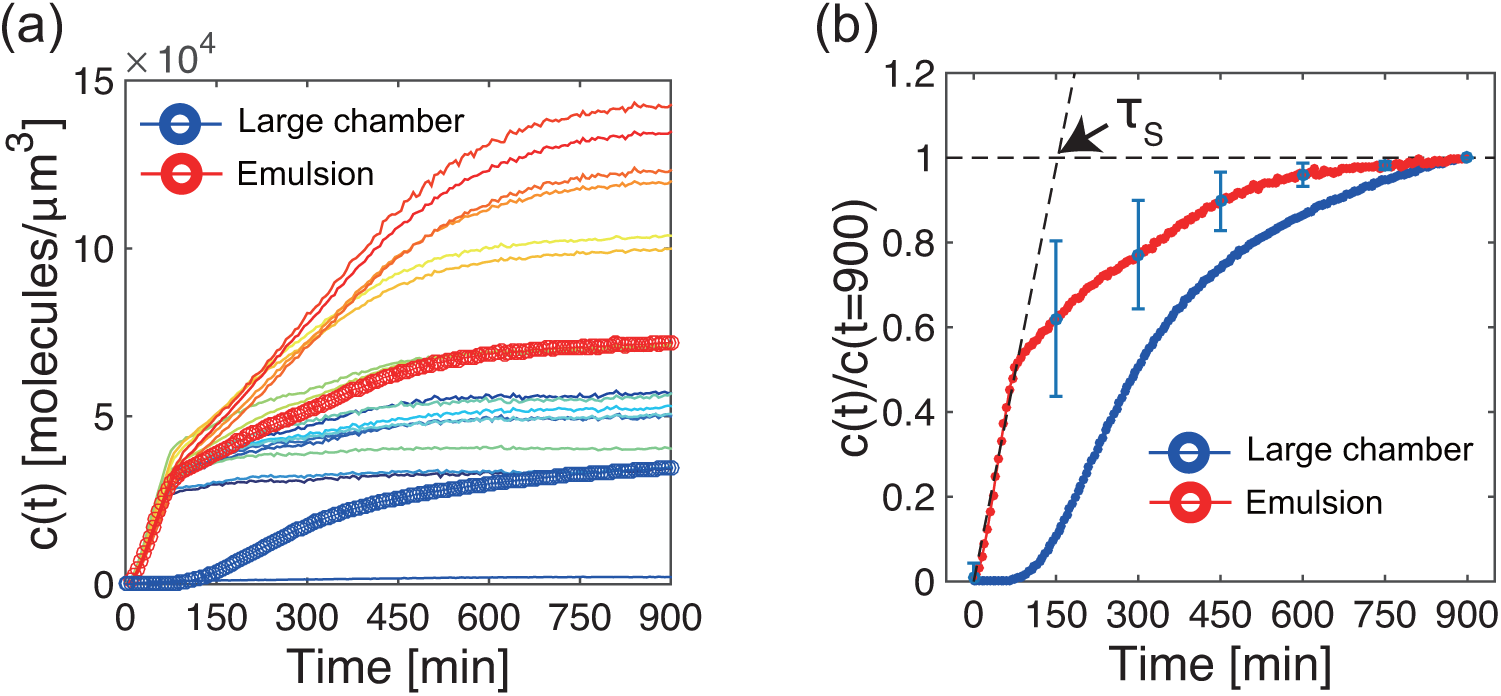
Time-course of protein synthesis (GFP concentration, *c*(*t*)) in various size of 18 droplets. The averaged kinetics over 18 droplets was plotted in red circle. Kinetics of deGFP production in CFES that was performed in a large frame-seal chamber was also shown in blue circle. (b) The time-courses of confined CFGE averaged over DOPC droplets (standard deviation is shown as blue vertical bars) and large chamber are normalized by the end-point GFP intensity. This curve was used to estimate the saturation time, *τ*_*s*_, in mathematical model

### Anomalous scaling of cell-free gene expression

The total fluorescent intensity *I*_*v*_ from Rhodamine-dextran (concentration *c*_*Rhd*_) scaled proportionally to the volume of each droplets *V*, so that normal scaling was observed as *I*_*v*_ = *c*_*Rhd*_*V* ∝ *R*^3^ (Fig. 3(a)). Interestingly, the signal *I*_*GFP*_, that was total fluorescent intensity of deGFP in each droplet, exhibited non-linear dependence to the volume marker *I*_*v*_ as shown in Fig. 3(a). The *I*_*GFP*_ for the expressed deGFP from TXTL is 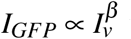 with *β* = 1.2, meaning that it is not linearly scaled with respect to the volume though *I*_*GFP*_ for purified deGFP is proportionally increased with the volume marker *I*_*v*_. This nonlinear dependence indicates that the CFGE within droplet is relatively suppressed if confined in smaller droplets.

**Figure 3.**
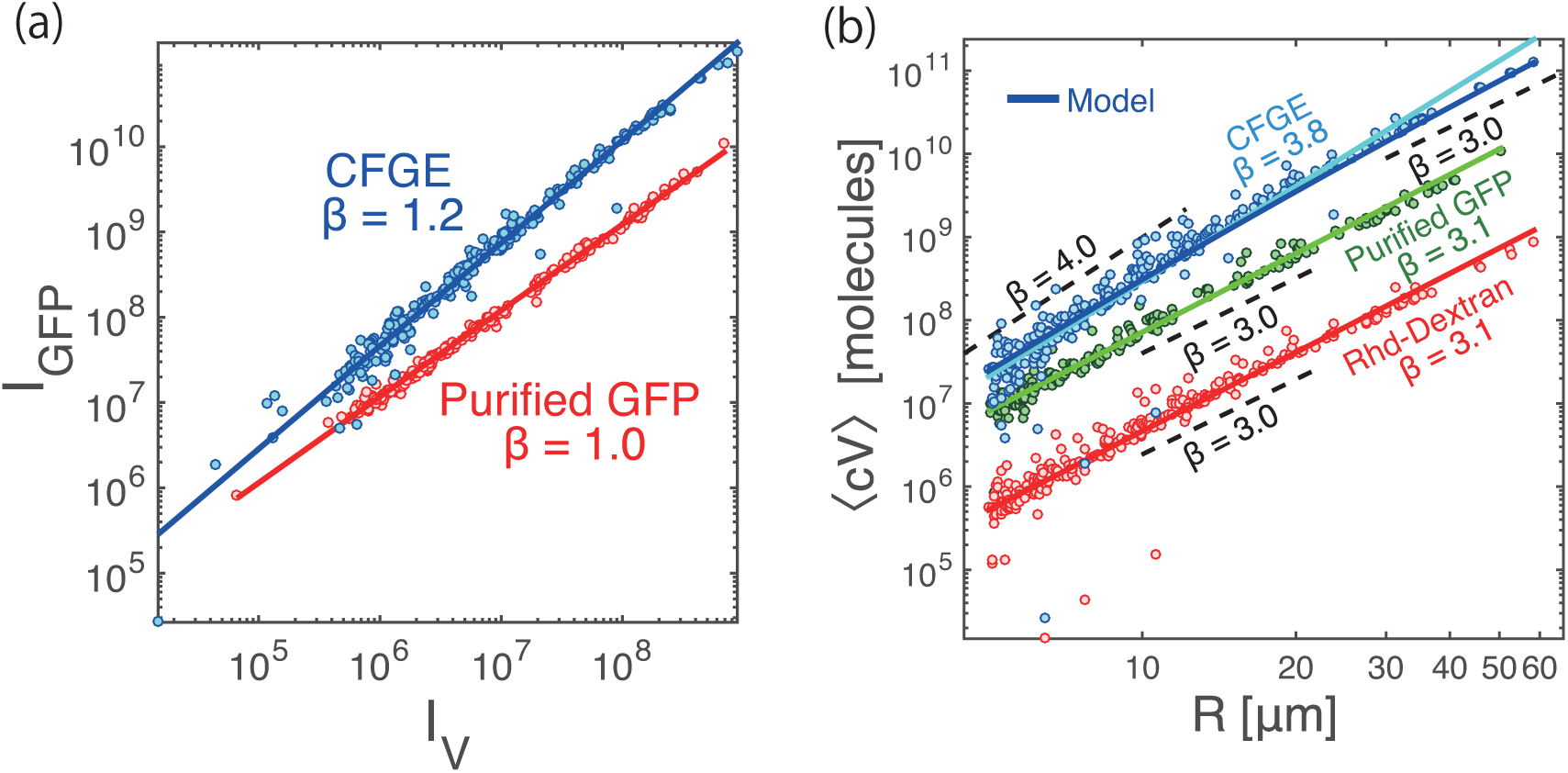
(a) deGFP expressed from TXTL extract (N=246) and purified deGFP (control, N=191) in each droplet. The horizontal axis represents the droplet size by using fluorescent reporter protein (Rhodamine-Dextran). (b) Log-log plots of the total intensity along droplet radius. Blue circles denote deGFP intensity resulting from CFGE in emulsion droplets. Green circles and red circles are the intensity of purified deGFP and marker protein, respectively. Blue circles are fitted to Eq. (7) using *k*_*R*_ = 3.21 × 10^−3^ [1/*μ*m^3^/sec], *k*_*on*_ = 1.0 [1/sec], *k*_*off*_ = 3.86 × 10^−3^[1/sec], *γ*_*R*_ = *γ*_*R*\*_ =5.00 [1/sec], *α* = 9.02 [1/sec], *α*\* = 1.96 × 10^−1^ [1/sec], *τ*_*s*_ = 153 [min], *λ* = 30 [nm] (blue solid line).

This result points out unexpected geometric dependence between outcome and input of CFGE under confinement.

To characterize this size-dependent CFGE, we further examined *I*_*GFP*_ in size-dependent manner. Fig. 3(b) shows *I*_*GFP*_ of expressed deGFP as the function of the radius of each droplet. We obtained a scaling function of *I*_*GFP*_ ∝ *R*^*β*^ with *β* = 3.8, suggesting that deGFP synthesized inside droplets follows an anomalous geometric scaling different from the ordinal *R*^3^ dependence. In contrast to expressed deGFP, *I*_*GFP*_ of DNA in emulsion exhibited *I*_*GFP*_ ∝ *R*^3.1^, therefore proportional to the volume (Fig. 3(b), red circle). Our next control experiments consisted of encapsulating purified deGFP protein inside the emulsion droplets. The *I*_*GFP*_ of purified deGFP still had *R*^3^ dependence, identical to the Rhodamine-dextran *I*_*v*_ ∝ *R*^3.1^. This control experiment indicated that the anomalous scaling *I*_*GFP*_ ∝ *R*^3.8^ is uniquely observed when deGFP is synthesized through TXTL inside the emulsion droplets.

### Mathematical model

The CFGE from the TXTL reactions has been described by Karzburn, et al^14^. Their coarse-grained mathematical model proposes a first order kinetics for mRNA degradation while the protein degradation follows a zeroth-order kinetics when deGFP is tagged for degradation. With no tag, deGFP cannot be degraded (infinite lifetime)^14,31^. We extend this model to be applicable for spherical confinement in order to elucidate the mechanism underlying in size-dependent CFGE.

Consider a cell-sized compartment of radius *R*, in which gene expression takes place and the products of transcription (TX) and translation (TL) cannot diffuse across the surface. Hereafter, we assume that protein synthesis in the vicinity of the droplet surface (thickness λ ≈ 30 nm) is regulated due to the activation/inactivation of TXTL onto the surface (Fig. 4(a)). We first give the mathematical model of gene expression under confinement. The time evolution of free mRNA (*r*), regulated mRNA (*r*) and produced protein (*c*) are described by the following rate equations:

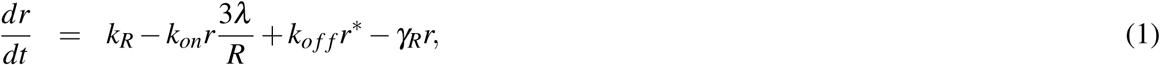

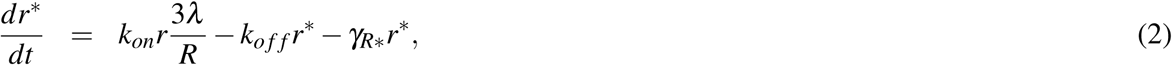

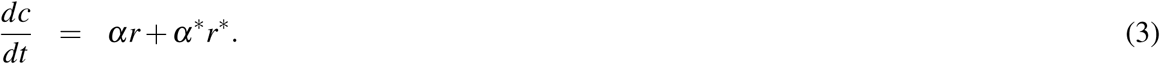

where the transcription rate of mRNA from DNA is *k*_*R*_, and the degradation rate of free mRNA and regulated mRNA are denoted as γ_*R*_ and γ_*R*\*_ respectively. mRNA produced from transcription is adsorbed to the thin surface layer of water-oil interface with the attachment rate *k*_*on*_ and the dissociation rate *k*_*off*_. The surface area to volume ratio of 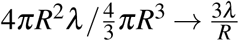 determines the capacity for mRNA in the surface layer, so that it leads the factor of 3*λ*/*R* in Eqs. (1) and (2).

**Figure 4.**
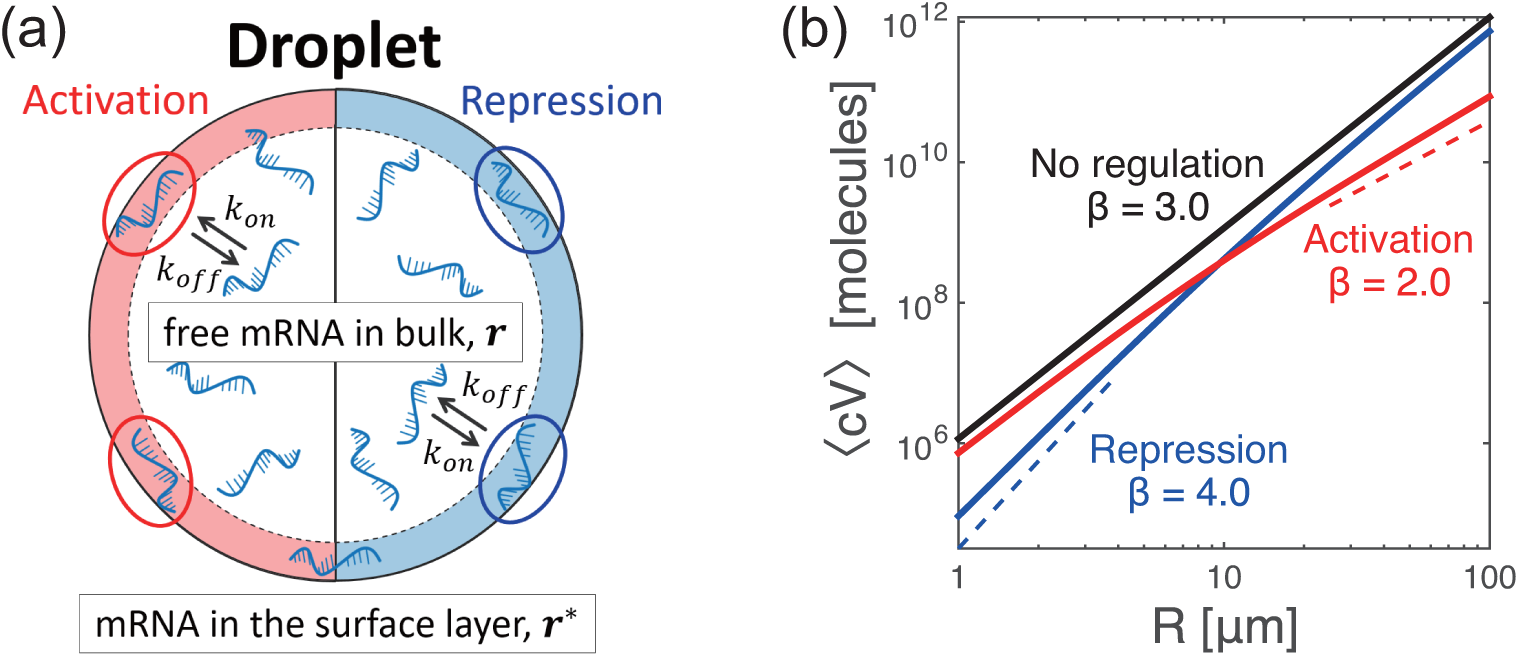
(a) Schematic illustration of the surface regulation model in a cell-sized compartment. (b) Theoretical analysis of anomalous scaling of produced protein ⟨*cV*⟩ in confined cell-free systems. Three types of regulation regime shows different size-dependent scaling. Repression (blue, *α* ≫ *α*\*), no regulation (black, *k*_*on*_ = 0), and activation (red, *α* ≪ *α*\*) correspond to *β* = 4.0, *β* = 3.0, and *β* = 2.0 for ⟨*cV*⟩ ∝ *R*^β^ respectively (broken lines are guide for the eyes.).

Synthesis of mRNAs rapidly reaches steady-state values ⟨*r*⟩ and ⟨*r*\*⟩ due to the short lifetime of mRNA molecules with respect to the observation time^14^. Steady-state expression is obtained by setting *dr / dt* = 0, *dr* / dt* = 0:

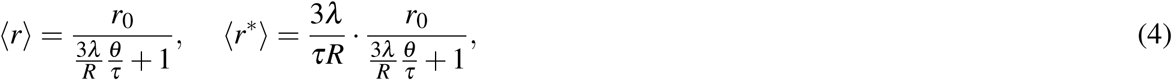

where 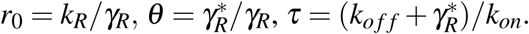 Multiply 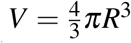 to Eq. (3) and substitute Eq. (4) into Eq. (3), then integrate it with the initial condition of *c*(0) = 0 and we obtain the time dependence of protein concentration:

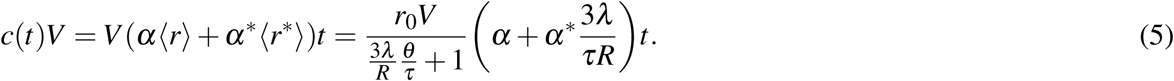

In our experimental setup, protein concentration finally saturates at a certain value due to the nutrients depletion^14^. Hence we determined the saturation time τ_*s*_ = 153 [min] from the intersection point between extrapolation of Eq. (5) and the experimentally observed plateau value (Fig. 2(b)). In the steady-state, therefore, Eq. (5) is rewritten as

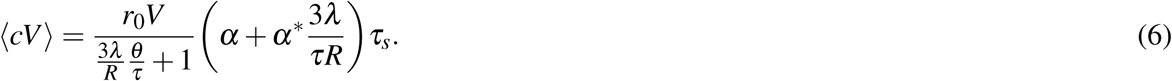

Eq. (6) represents general geometric scaling of CFGE under confinement, so that we next examine the exponents of size-dependent scaling under two kinds of gene regulation, namely either surface-induced repression (*α* ≫ *α*\*) or surface-induced activation (*α* ≪ *α*\*). Firstly, for the surface-induced repression of TX/TL, given that *α* ≫ *α*\* yields as the repressed protein synthesis, Eq. (6) becomes

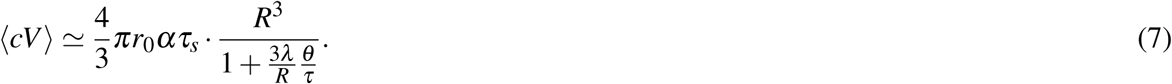

To find simple scaling relation, one expects that the following relation holds: *θ* ≫ *τ*, i.e. 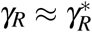 and *k*_*on*_ ≫ *k*_*off*_ + *γ*_*R*_, meaning that the translation of mRNA suppresses at the surface until mRNA is degraded.

This approximation gives quartic dependence of CFGE on *R*: ⟨*cV*⟩ ∝ *R*^4^ which tells that for small droplets translation on the surface layer is effectively suppressed, while the proportion of surface-induced repression becomes negligible for large droplet and approaches to *R*^3^. We found that this size-dependent scaling Eq. (7) agrees well with the experimental result shown in Fig. 3(b), suggesting *I*_*GFP*_ ∝ *R*^4^ for CFGE in DOPC droplets. We thus conclude that TXTL reaction in droplets belongs to the class of surface-induced repression.

In contrast, surface-induced activation, when mRNA in the surface layer is much activated rather than bulk (*α* ≪ *α*\*), Eq. (5) becomes

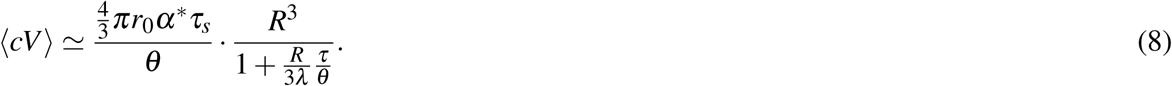

For large droplets, Eq. (8) becomes ⟨*cV*⟩ ∝ *R*^2^, whereas size-dependence becomes *R*^3^ for small droplets. This kind of size-dependence was not observed in our experiment. These two anomalous scalings of ⟨*cV*⟩ are shown in Fig. 4(b). Simple theoretical model thus proposes how the surface alters the size-dependent scaling of protein synthesis.

## Discussion

We have demonstrated the existence of geometric regulation of CFGE and its anomalous scaling in a cell-sized compartment. Our study shows how important are the surface properties in changing biochemical reactions kinetics. Theoretical model hypothesizes that TXTL is suppressed in a thin layer close to the membrane surface and that it alters the geometric scaling law between synthesized protein and the size of compartment. CFGE in water-in-oil droplets follows a surface repression regime, consequently the end-point protein concentration inside droplets varies with droplet size in emulsion. Non-catalytic compartmentalization thus represses. Although it has been known that electrostatic interactions make polyelectrolytes of DNA localized by electro-static interaction with charged membrane^32^, our finding points out that the geometric regulation of CFGE occurs even at the interface abundant of neutral lipids, implying that steric interaction alone in thin surface layer (typically thickness is few tens of nm) gives rise to control CFGE. Although such surface-induced repression of TXTL in a cell-sized droplet is evident in emulsified droplet, the CFGE from PURE system in liposomes^33,34^ appears to show normal scaling behavior of *I*_*GFP*_ ∝ *R*^3^. The PURE system has better stability against mRNA degradation rather extract-based system^35,36^, the approximation *k*_*on*_ ≫ *k*_*off*_ + *γ*_*R*_ is reasonable as well. Such discrepancy may result from the difference of physical nature of CFES: TXTL extract in this study is at rather crowded situation due to the presence of macromolecules such as few percent of polyethylene glycol with abundant protein machinery. On the other hand, liposomes containing PURE system without any additional macromolecules will be difficult to support the surface layer, implying that the rate of translation close to the membrane surface could be comparable to that in bulk (*α* ≈ *α*\*). Cell-sized surface gives rise to the anomalous scaling of CFGE though the experimental verification of surface-induced activation, *I*_*GFP*_ ∝ *R*^2^, will be addressed in future work.

The anomalous scaling of gene expression points out that confinement with stable surface can alter the protein synthesis even in simple cell-free systems. This fact in turn brings practical strategy to design how the functional surface of artificial bioreactors capable of rational control of gene expression. For small compartment as small as bacteria, because the surface area to volume ratio is large, the regulation of CFGE associated with membrane becomes more effective. For surface-repression strategy, confinement in small droplets may be useful to save chemical resources for protein synthesis. When the droplets grow or coalesce until their size becomes sufficiently large^19,37^, the droplets will recover normal size-dependence of *R*^3^. In contrast, to suppress the protein synthesis at larger droplets, surface-induced activation is favorable due to another anomalous scaling of *R*^2^. Thus, the sign of surface-induced regulation is important to build functional interface in confined cell-free bioreactors. Further investigation of CFGE in a confined space will advance a versatile miniaturized factory of protein synthesis in cell-free bioreactors^9,15,29,30^, and in turn will bring better understanding of macroscopic geometry and microscopic enzymatic kinetics in a cell-sized space^10,18^.

## Methods

### Image analysis

Quantitative image analysis was done using custom code written in MATLAB^31^. Specifically, the bright field image taken at end point was processed through binarization and in turn the center of mass of each droplets was detected. To obtain the radius of droplets of *R*, the area of each droplets were extracted and then assumed to *πR*^2^. To determine *R* with better precision and reliability, we performed two filtering processes. When the water-in-oil droplets spread onto the substrate surface due to wetting or when the coalescence of multiple droplets occurred, such droplets exhibited deformed ellipsoid-like shape. Because such asymmetry causes potential difficulties in subsequent data analysis, we excluded the debris according to two filtering methods. First filtering method was used to figure out whether detected objects were spherical droplets or deformed one. We defined a metric, *M*, to evaluate the roundness of an object: *M* = 4*π*  *S=L*^2^, where *S* and *L* is the area and the perimeter of the object, respectively. This metric becomes 1 for a spherical object while less than 1 for any other shape. Its precision was adjusted with an appropriate threshold and we employed threshold *M* = 0.75 in this study. Second filtering method is to remove small debris of micelles out of detected objects. The detected objects whose size was less than 78.6 *μ*m^2^ were excluded from subsequent analysis. This filtering process sets the minimum limit in droplet radius at 5 *μ*m. In addition, background intensity was subtracted from each pictures, where background region was chosen in a way to be sure no background pixels were in droplets.

### Protein purification

Purified deGFP with histidine-tag at C-terminal end was used in one of control experiments. The expression vector of deGFP was constructed by using pET28b(+) plasmid DNA with conventional cloning method. The deGFP was expressed in *Escherichia coli* strain of BL21(DE3)pLysS (TaKaRa) with selective antibiotics and then purified through Ni-NTA column (QIAGEN). After measuring the concentration of purified deGFP by both Bradford assay and gel-electrophoresis, the calibration curve of fluorescent intensity of deGFP in fluorescent microscopy and protein concentration was obtained.

## Acknowledgements

We thank Z. Izri for fruitful discussion. This work was supported by Human Frontier Science Program Research Grant (RGP0037/2015) and JSPS KAKENHI (“Synergy of Structure and Fluctuation” 17H00225, “Hadean Bioscience” 17H00245, and Grant-in-Aid for Scientific Research (B) 17KT0025) from MEXT.

## Author contributions statement

R.S. and Y.T.M designed the experiments, V.N. prepared the cell-free extract, R.S. conducted the experiments and analyzed the results. R.S. and Y.T.M. constructed mathematical model. All authors wrote and reviewed the manuscript.

## Supplemental information for Anomalous Scaling of Gene Expression in Confined Cell-Free Reactions

### Experimental details

#### Image analysis

**Figure 5.**
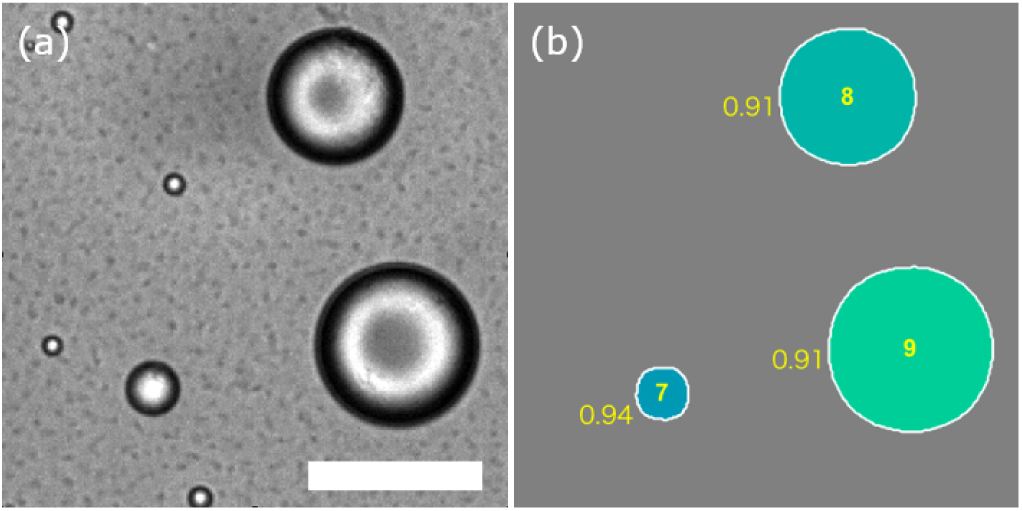
(a) GFP fluorescence within droplets (bar: 40 *μ*m). (b) Droplets are labeled by the numbers in the center of each droplet. Three-digit numbers outside of the droplets denote the metric of each droplet. The small debris and coalesced droplets were excluded after filtering methods.

#### Protein degradation in confined cell free extract

It has been shown that protein degradation in cell free extract is zeroth order^14^. As for GFP reporter protein used in this study, protein degradation is assumed to be negligible owing to the absence of ssrA peptide tag responsible for protein degradation. To test whether the degradation of GFP is negligibly small, we measured the decay of fluorescent intensity of purified deGFP in the TXTL extract within DOPC droplets. The time course of fluorescent intensity is shown in Fig. 6(a), where data curve is normalized by the end-point (900 min) thereafter averaged over all droplets (red curve) and standard deviation is shown (blue vertical bars). Intensity fluctuation is observed in the time course due to the slight moving of each droplet, however, there is no considerable intensity decrement. In addition, we also tested the decay of purified GFP protein in the phosphate buffer solution within DOPC droplets but its considerable reduction was negligible (Fig. 6(b)). We thus conclude that there is no active degradation in TXTL extract as well in this study.

**Figure 6.**
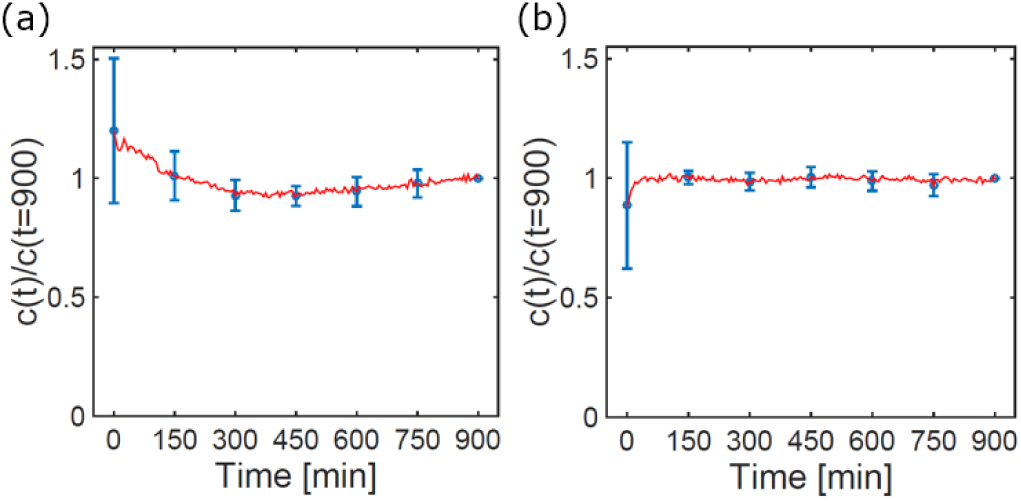
(a) Time course of normalized GFP intensity in water-in-oil droplets comprising of TXTL extract and purified deGFP, (b) comprising of PBS (Phosphate-buffered saline) and purified deGFP. The red curve is average for over all droplets ((a): *N* = 29, (b): *N* = 24) and standard deviation is shown as blue vertical bars.

